# Social isolation impacts late-life socio-cognitive decline in APP/PS1 mice

**DOI:** 10.1101/777441

**Authors:** Eva Rens, Rudi D’Hooge, Ann Van der Jeugd

**Affiliations:** Laboratory for Biological Psychology, Tiensestraat 102, KU Leuven, Leuven, Belgium

**Keywords:** aging, dementia, cognition, sociability, animal models

## Abstract

In this study the effects of social isolation (SI) were investigated in APP/PS1 mice. It was found that SI during adolescence has an impact on anxiogenic behaviour, such that isolated animals tend to explore a threatening environment less than non-isolated animals as assessed with the EPM test, and that this holds for both AD and non-AD mice. While no evidence was found for any differences in short-term memory as assessed by the Y-maze, long-term memory seemed to be affected in a context-dependent manner. Object memory as assessed with the NOR test was affected in APP/PS1 mice compared to WT mice, but this deficit was not induced or influenced by SI. When it comes to social recognition memory however, we found that SI exacerbated the social memory deficit in AD mice, and even induced a deficit in WTs. Associative fear memory as assessed with the PA test suggested that WTs perform better when group housed, and APP/PS1 mice better when socially isolated. The link between isolation and AD, or cognition in general, may be more complex than initially thought. The effect of isolation may not be the same for AD versus non-AD subjects.

## Introduction

The past years, researchers and health practitioners have identified social isolation (SI) as one of the risk factors for developing neuropsychiatric disorders. SI, both objective and perceived, leads to several detrimental health conditions through three main pathways: via behavioural, psychological, and physiological health (Berkman, Glass, Brisette & Seeman, 2000). At the behavioural level, an older adult’s social network can impact health through provision of social and instrumental support, through social influence and through social engagement (Cacioppo & Hawkley, 2003). SI also acts through psychological pathways. Isolated elderly are at an increased risk for depressed mood and hopelessness (Cacioppo & Cacioppo, 2013; Golden et al., 2009), and eventually, suicide attempts (Lebret, Perret-Vaille, Mulliez, Gerbaud & Jalenques, 2006). Moreover, SI in older adults is associated with poorer overall cognitive functioning and even increases the risk on dementia (Cacioppo & Cacioppo, 2013; DiNapoli, Wu & Scogin, 2014; Fratiglioni, Wang, Ericsson, Maytan & Winblad, 2000). Finally, also physiological processes have been linked with SI. A meta-analysis focusing on the cardiovascular, endocrine and immune systems as potential physiological mechanisms found that a lack of social support appeared most reliable on blood pressure, catecholamines and aspects of both the cellular and humoral immune response (Uchino, Cacioppo & Kiecolt-Glaser, 1996). Both perceived and objective SI are associated with increased all-cause mortality (Steptoe, Shankar, Demakakos & Wardle, 2013).

AD is a neurodegenerative disorder characterised by a chronic and progressive course (Burns & Iliffe, 2009). Two types of brain lesions are idiosyncratic for AD: amyloid deposits (often called plaques) resulting from the aggregation of amyloid-β (Aβ) peptide, and neurofibrillary tangles of hyperphosphorylated pathologic tau proteins (Vinters, 2015). Aβ plaques are extracellular whereas NFTs develop within neurons. Also characteristic of AD is the enlargement of ventricles and shrinkage of the subcortical white matter (Vinters, 2015).

SI increases the risk for neuropsychological disorders and mortality in elderly (Friedler et al., 2015; Gilman et al., 2015; Jiang, Cowell, & Nakazawa, 2013; O’Keefe et al., 2014). Research suggests that SI is a potential risk factor for age-dependent cognitive impairments, dementia and more specifically AD (Leser & Wagner, 2015). SI is associated with cognitive decline and a twofold risk on the development of AD-like dementia (Wilson et al., 2007). It was recently found that SI and loneliness had a higher prevalence in AD patients compared to a group of healthy controls (El Haj et al., 2016). SI is also found to be linked with cortical amyloid burden in healthy adults, further suggesting that SI could be a manifestation relevant to preclinical AD (Donovan et al., 2016).

Animal studies provide more standardised evidence of the role of objective SI as a risk factor of AD. Complete SI was found to induce neuronal degeneration in both AD and non-AD rats, and enhance the severity of AD development in isolated AD rats compared to socialized AD rats, for example by increasing Aβ deposition (Ali, Khalil, Elariny & Abu-Elfotuh, 2017). SI was shown to exacerbate the impairment of spatial working memory and the accumulation of Aβ42 in adult APP/PS1 mice (Huang, Liang, Ke, Chang & Hsieh-Li, 2011). In seventeen-month-old APP/PS1 mice these effects are even more pronounced, and isolation housing was found to decrease learning, memory functions and exploratory behaviour, and exacerbate AD pathology such as accumulation of Aβ and hippocampal volume loss (Huang et al., 2015). However, no study looked at the effects of SI on emotionality and social behaviours in APP/PS1 mice.

In order to examine the link between SI and AD in more detail, in the present study, we compared the phenotype of isolated mice to that of group housed mice in both healthy aging as well as AD-like pathology (APP/PS1 transgenic mice) in a series of behavioural tests probing for several fields of cognition, sociability, emotionality and exploration. Since perceived and objective SI are highly prevalent in our society, especially in young adults and the very old, this may have implications for developing or accelerating AD pathology. Insight in how environmental conditions can influence cognition and disease processes is important for optimising interventions. For example, individuals at risk for SI can be targeted for support and this may have the potential to reduce the risk on developing AD. AD is a global public health priority given its rising prevalence and enormous cost on society, so we believe that every step with the potential of reducing AD is worth taking. Especially since treatment is not possible yet, prevention should be at least as important to focus on.

## Materials and Methods

### Animals and housing conditions

A total of 64 mice was used, 50 APP/PS1 mice, also 14 healthy wild-type (WT). APP/PS1 mice were used as a disease model of AD. APP/PS1 mice contain human transgenes for both APP containing the Swedish mutation causing FAD, and for PSEN1 containing an L166P mutation, both under the control of the Thy1 promotor on a C57BL/6J background (Radde et al., 2006). APP/PS1 mice contain three times more human APP than endogenous APP (Maia et al., 2013). Deposition of amyloid plaques starts in the neocortex at approximately six weeks of age, in the hippocampus at three to four months (Radde et al., 2006). WT mice were used as control.

About half of the mice were group housed (GH) with same-sex congeners, the other half were single housed and thus socially isolated (SI) for a month prior to and for an additional month during experiments. All animals had continuous access to water and food and were kept on a 12 h light/dark cycle. Cage beddings consisted of wood shavings and environmental enrichment was provided with an empty roll of toilet paper and paper tissue that served as nesting material. The week prior the behavioural testing, the mice were gently handled for 5 min per day in order to become familiarised to the experimenter. Behavioural testing started at 15 months of age.

#### Procedure

All tests were performed according to the standard operating procedures of the Laboratory of Biological Psychology, KU Leuven. The order of the tests is based on an implied ranking going from least to most invasive and stressful. Elevated plus maze, open field, novel object recognition, social preference social novelty, passive avoidance and forced swim test.

## Exploration, anxiety and depression-like behaviour

### Elevated plus maze

The apparatus consisted of a plus-shaped maze with two open and two closed arms covered with a dark lid, both 5 cm wide and 20 cm long. The mouse was put at the end of a closed arm and its activity was then recorded for 5 min without habituation period. Visits to the arms were registered by four infrared beams (one for each arm entry) and sent to an activity logger. Time spent in the open arms was calculated as a measure for anxiety.

### Open Field

Open Field (OF) exploration was assessed by introducing the mouse individually in the corner of a novel and brightly lit (650 lux) plexiglas box with non-transparent walls consisting of a 50 x 50 cm^2^ square arena with a height of 30 cm. The box was placed in a closet and the doors were shut to ensure minimal influence of external stimuli. A camera located above the box tracked the activity of the mouse and transmitted information to a computer equipped with ANY-maze™ (Stoelting Co., IL, USA) behavioural tracking software. The animal was tracked for 10 min. Prior to the start of the OF test, mice were dark habituated for 30 min in a closed cupboard (other from the OF one). The centre of the OF was defined as the inner circle with a diameter of 30 cm, the periphery was defined as 0-5 cm from the walls. After each trial, the arena was thoroughly cleaned with ethanol in order to eliminate scent trails. The mean speed and total distance covered were assessed, which are expressed in cm/s and cm respectively and are measures of general locomotor activity. The variable of interest for exploratory activity is the time spent in the centre.

### Forced swim test

Mice were placed in a cylindrical glass tank filled with 6 l water at 25 °C. The animals were considered immobile when they did not attempt an escape the tank, but just floated with only minor movements necessary to keep their head above water. Behaviour was video-taped and immobility was visually judged and recorded manually by two observers during 5 min. Two variables were of interest: the total time the mouse was immobile during the six min test and the latency until first immobility.

## Working memory

### Y-maze

The Y-maze consisted of one base arm and two target arms each 35 cm long, 5 cm wide and at 120° from each other. A forced-choice spontaneous alternation paradigm was used. In the acquisition phase, the mouse could freely explore the maze for 2 min, but one target arm was closed off with a small door. The test phase with both arms accessible took place after an inter-trial interval of 2 min. The left and the right target arm were blocked an equally amount of times during the acquisition phases throughout the experiment in order to control for side preference effects. After each trial, the maze was thoroughly cleaned with ethanol. As a measure of short-term memory, the first arm entered in the test phase was used.

## Long-term memory

### Novel object recognition

The NOR test took place in the same Plexiglas OF box, 400 lux lit, and the animal’s activity was tracked using ANY-maze™ (Stoelting Co., IL, USA). In the acquisition phase, two identical objects were placed near two corners of the box, and the mouse could freely explore for 5 min. The test phase took place after a delay of 24 h and lasted 5 min. Randomization ensured that for half of the animals the left object was novel, and for the other half the right object novel. Both the arena and the objects were thoroughly cleaned with ethanol after each mouse. Time spent exploring the novel object were recorded with the tracking apparatus. The discrimination ratio (DR) was calculated for time visiting the novel object by dividing the time at the novel object by the total amount of time spent with both the object.

### Social recognition memory

The SPSN test took place in the Three-chamber box, 400 lux lit, and the animal’s activity was tracked using ANY-maze™ (Stoelting Co., IL, USA). The two outer chambers (w × d × h: 29 × 28 × 30 cm) were connected with the center by gates in transparent walls (w × h: 6 × 8 cm). A cylindrical wire cup (h × Ø: 11 × 12 cm) was provided in the center of each outer chamber. Here, mice could be placed as stimuli for the test subjects. During the habituation phase that lasted 5 min, the middle compartment served as an empty habituation zone with closed gates. During the second phase, a stranger mouse was added in the left or right chamber. In this phase, sociability was measured for 10 min. During this social preference test, the mouse could choose between spending time at the empty side, the habituated center or at the stranger’s side (the latter represents social exploration). During the final phase, preference for social novelty was measured for 10 min. In this social recognition memory test, the mouse could choose to prefer the company of the first or newly introduced stranger. During the experiment, the stranger mice had their own round wire cage that made auditory, tactile, olfactory and visual contact possible. At every trial, these cages were placed at the same spots in the boxes. For ethical reasons, their stay in the cages was as short as possible. When analyzing social preference, the parameter of interest is DR of time to the proximity of the stranger’s compartment in comparison to the empty compartment. When analyzing the social novelty preference, the DR of new stranger preference above the old stranger is of interest. The discrimination ratio (DR) was calculated for time visiting the novel mouse by dividing the time at the novel stranger by the total amount of time spent with both strangers.

### Fear memory

The passive avoidance test was used to probe associative fear-related long-term memory. The apparatus consisted of a small, brightly (650 lux) illuminated Plexiglas box connected via a gate to a dark chamber fitted with a grid floor connected to a constant current shocker. A step-through paradigm was used. Animals were adapted to the dark 30 min prior to the testing to ensure aversion of light conditions. In the acquisition phase, the animal is placed in the light box and receives access to the dark area after 10 s. A light foot shock of 0.3 mA is given for 2 s when the animal enters the dark chamber (training latency). The retention test took place after a delay of 24 h. The test latency to enter the dark was measured and a cut-off time of 3 min was used. No shock was given this time.

## Results

### SI increased anxiogenic behaviours in mice, but did not affect general and social exploration behaviours, nor depression-like behaviours

One mouse was excluded from the analysis. No significant main effect of genotype was found for the proportion spent in the open arms of the EPM (*F*(1,55) = .865, *p* = .365). A strong main effect of housing condition was found, SI mice significantly explored the open arms less than the GH group (*F*(1,55) = 8.467, *p* = .005, see Figure 1). No effect of gender was found in the EPM (*F*(1,55) = .012, *p* = .912). Interaction effects were also absent.

**Figure 1.**
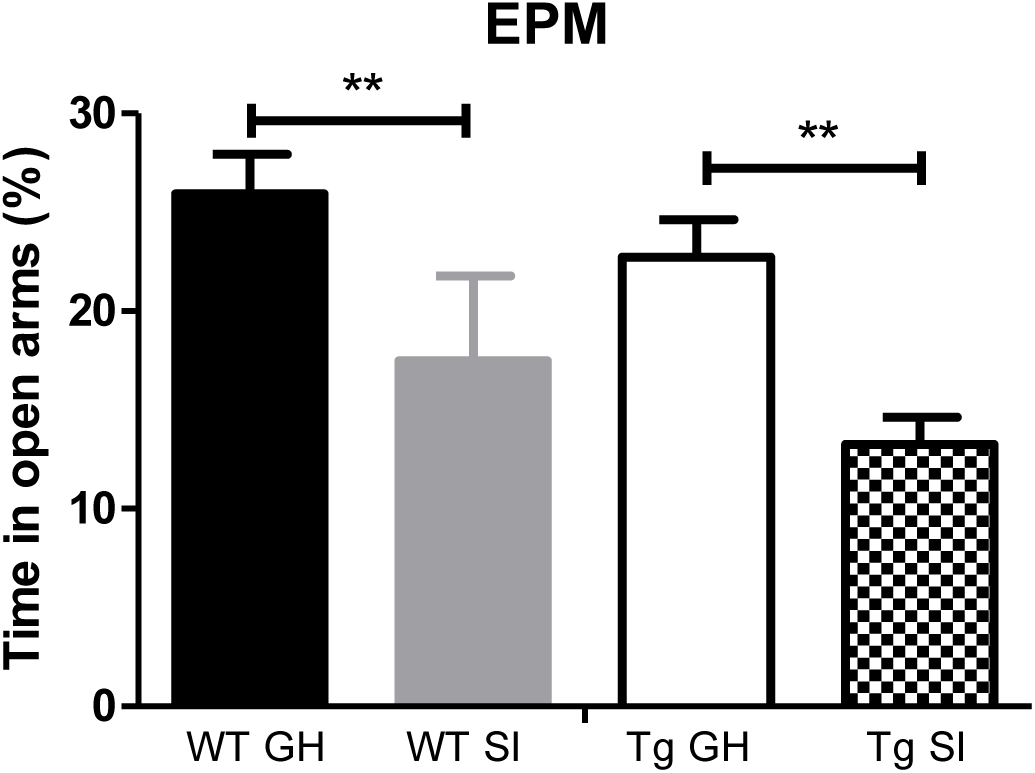
The mean proportion spent in the open arms in the elevated plus maze. Social isolation leads to a decrease in exploration of the open arms of the elevated plus maze, regardless of the genotype status or gender (** *p* < .01).

One mouse was excluded because it remained immobile for 95% of the time in the OF. Mean speed and total distance covered were investigated to assess general locomotor abilities. No effects were found for speed and total distance (respectively *F*(1,55) = 3.59, *p* = .063; and *F*(1,55) = 3.614, *p* = .068).

A main effect of genotype was found for time spent in the centre, APP/PS1 mice spent less time in the centre then WT mice (*F*(1,55) = 15.092, *p* < 0.001). The effect of housing condition was not significant on time spent in centre (*F*(1,56) = 1.669, *p* = .045, *η*^2^ = .071, see Figure 2. No effect of gender was found, *F*(1,56) = 2.548, *p* = .116, *η*^2^ = .044, and also interaction effects were absent.

**Figure 2.**
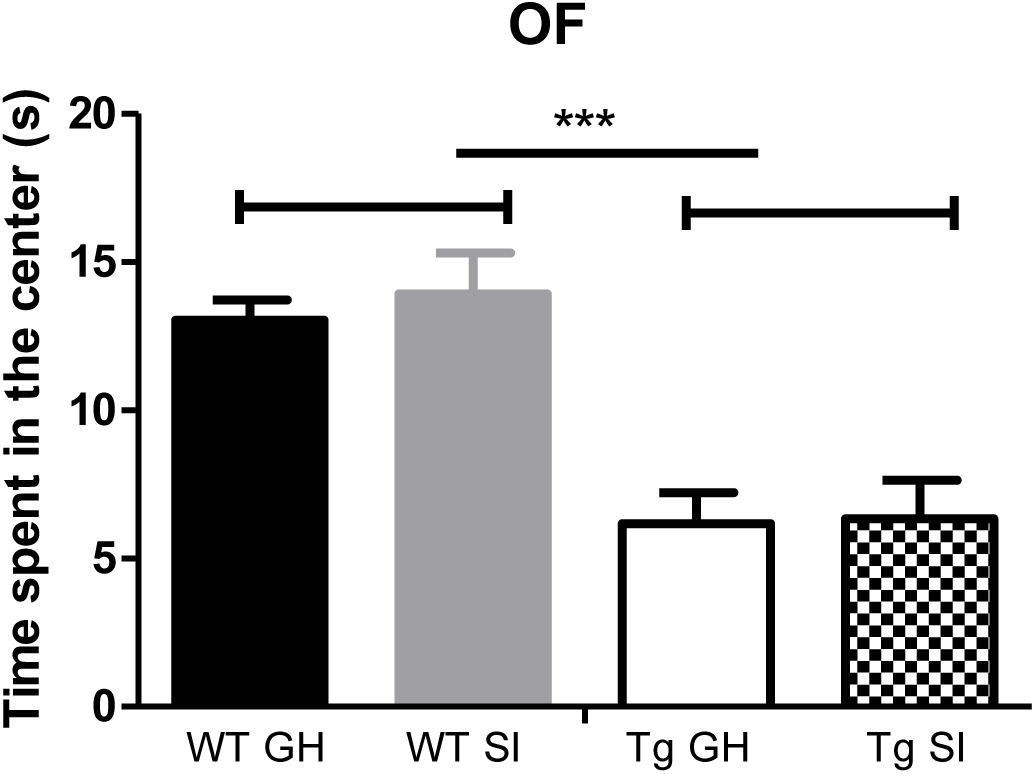
Time spent in the centre in the open field test. Social isolation led to a decrease in exploration of the centre of an open field arena, regardless of the genotype status or gender (*** *p* < .001).

The amount of social exploration, operated as the time spent close to the unfamiliar congener in phase 1, differed significantly between genotypes, *F*(1, 55) = 8.501, *p* = .005, see Fig 3, indicating that the APP/PS1 group explored their congener less than the WT group.

**Figure 3.**
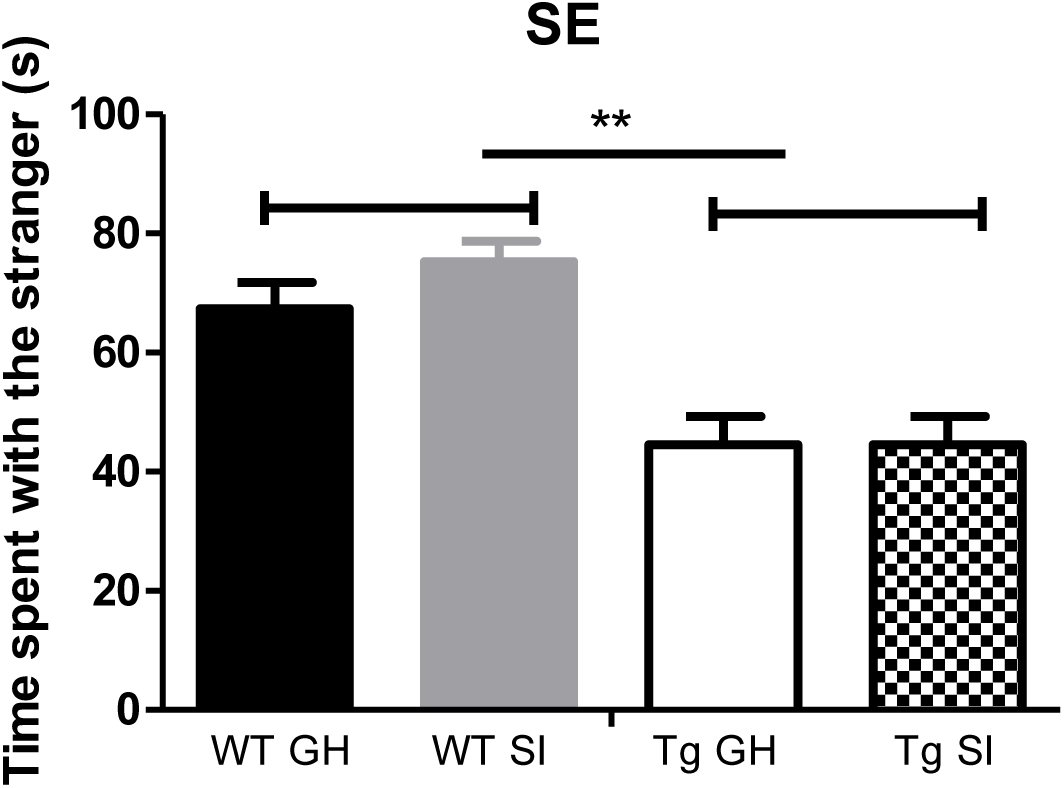
Time spent near the stranger in the social exploration test revealed that APP/PS1 mice spent less time exploring the stranger mice compared to WT mice (** *p* < .005).

Somewhat surprisingly, no main effect of housing condition was found for time spent close to the congener, *F*(1, 55) = 1.927, *p* = .171. Also a significant effect of gender was absent in time spent near the congener, *F*(1,55) = .363, *p* = .548. No interaction effects were present in the SE test. For the social novelty test following the social preference phase, please see social memory paragraph.

For the variable ‘first time immobile’, there were no main effects of genotype, *F*(1,56) = 1.024, *p* = .316, housing condition, *F*(1,56) = .550, *p* = .0461, or gender, *F*(1,56) = .039, *p* = .845, data not represented. Also interaction effects were absent.

For the variable ‘total time immobile’, APP/PS1 mice were significantly more immobile than WT mice, *F*(1,56) = 5.936, *p* = .018, see Figure 4. This was the only main effect present, as housing condition, *F*(1,56) = 2.259, *p* = .138, and gender, *F*(1,56) = .413, *p* = .523, revealed no effects. Interaction effects were again absent.

**Figure 4.**
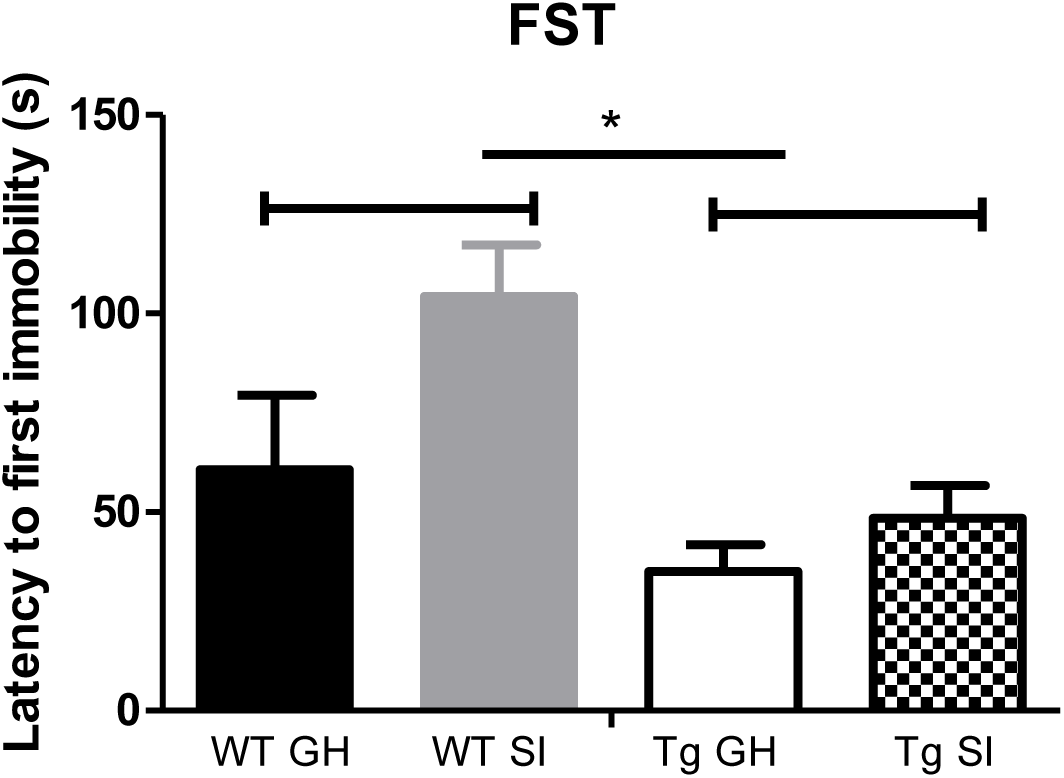
Latency to first immobility in the forced swim test revealed that APP/PS1 mice more rapidly displayed defeat behaviours compared to WT mice (* *p* < .01). Isolation did not affect this depression-like behaviour.

### SI did not affect working memory

Two mice were excluded because they did not explore the maze in the first phase. For the proportion of visits to the novel arm, no significant main effect was found of genotype (*F*(1,54) = 2.591, *p* = .113), housing condition (*F*(1,54) = .006, *p* = .941), or gender (*F*(1,54) = .099, *p* = .754; Fig 5). Also interaction effects were not found. With 52% (*SD* = .505) correct visits of the APP/PS1 group compared to 71.429% (*SD* = .469) of the WT group, the APP/PS1 group has a stronger *trend* towards lower memory performance.

**Figure 5.**
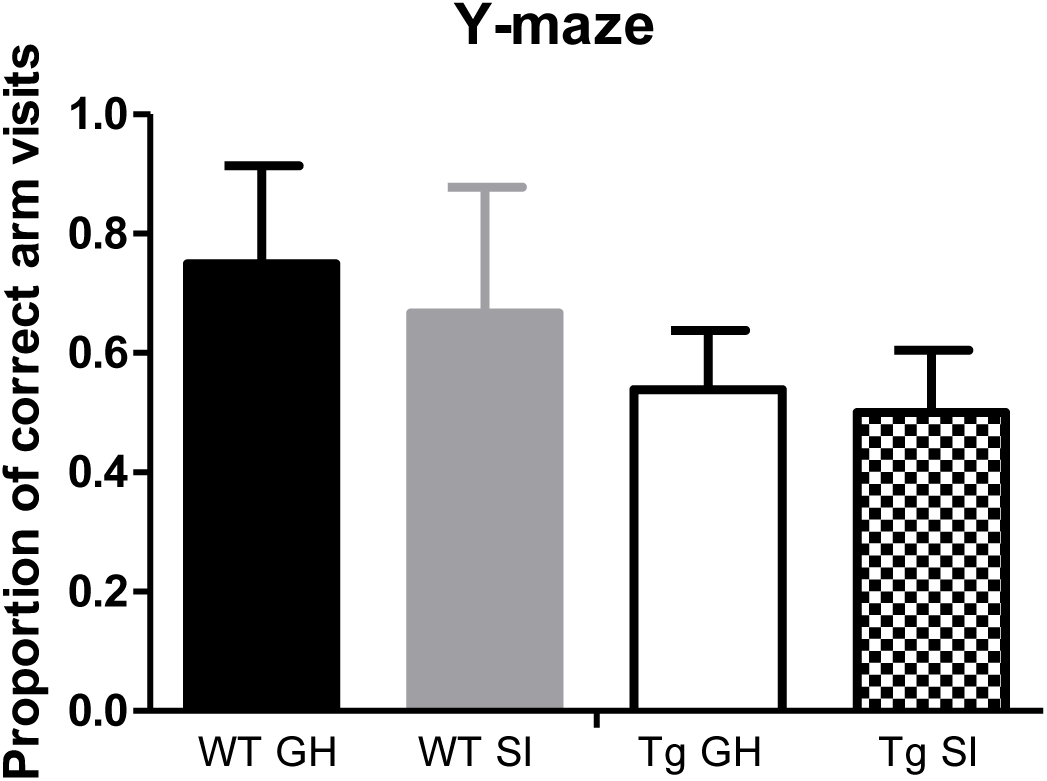
Proportion of visits to the novel arm in the Y-maze was not different after social isolation (all *p* > .05).

### SI did not influence novel object recognition memory, but exacerbated social retention memory deficits in APP/PS1 mice

Two mice (APP, GH, male; APP, SI, female) explored only one object during the training phase. One mouse (WT, GH, female) never explored the novel object during the test phase. For the first object visited, no significant main effects were found of genotype, *F*(1,55) = .063, *p* = .802, housing condition, *F*(1,55) = .732, *p* = .396, and of gender, *F*(1,55) = .804, *p* = .374. Interaction effects were absent as well.

When looking at the DR of time spent with the novel object, no main effect of housing condition was found, *F*(1,53) = .304, *p* = .584. The effect of genotype was significant, *F*(1,53) = 4.151, *p* = 0.047, and no main effect of gender *F*(1,53) = .521, *p* = .474, and interaction effects were found (Figure 6).

**Figure 6.**
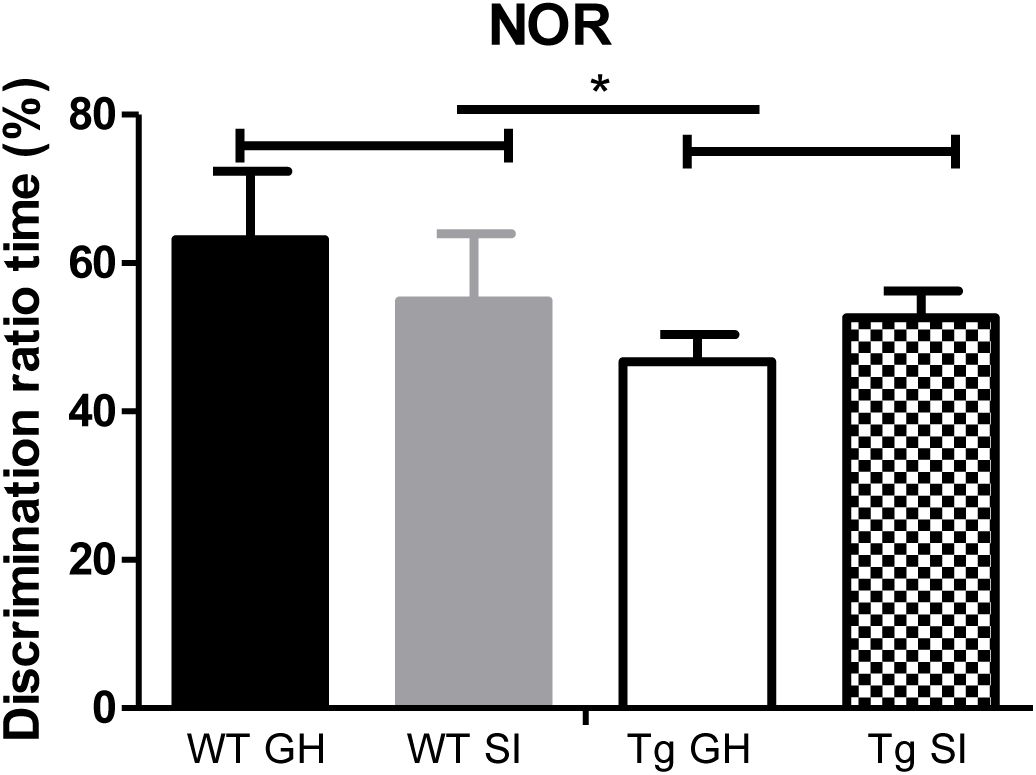
Discrimination rate (DR) of time spent exploring the novel object revealed that APP/PS1 mice had poorer object retention memory then their healthy conspecifics (\**p* < 0.05).

One mouse (WT, GH, female) never explored the stranger mouse during the test phase. When looking at the DR of time spent with the novel object, an effect of genotype was found, *F*(1,55) = 5.507, *p* = 0.023, meaning that APP/PS1 mice had poorer social memory compared to WT mice. Remarkably, a main effect of housing condition was also found, *F*(1,55) = 6.354, *p* = .015, signifying that isolated WT and APP/PS1 mice suffered from poorer memory then their group-housed conspecifics. No main effect of gender *F*(1,55) = .001, *p* = .977, and also surprisingly no interaction effects (especially the absence of Genotype x Housing Condition) were found (Figure 7).

**Figure 7.**
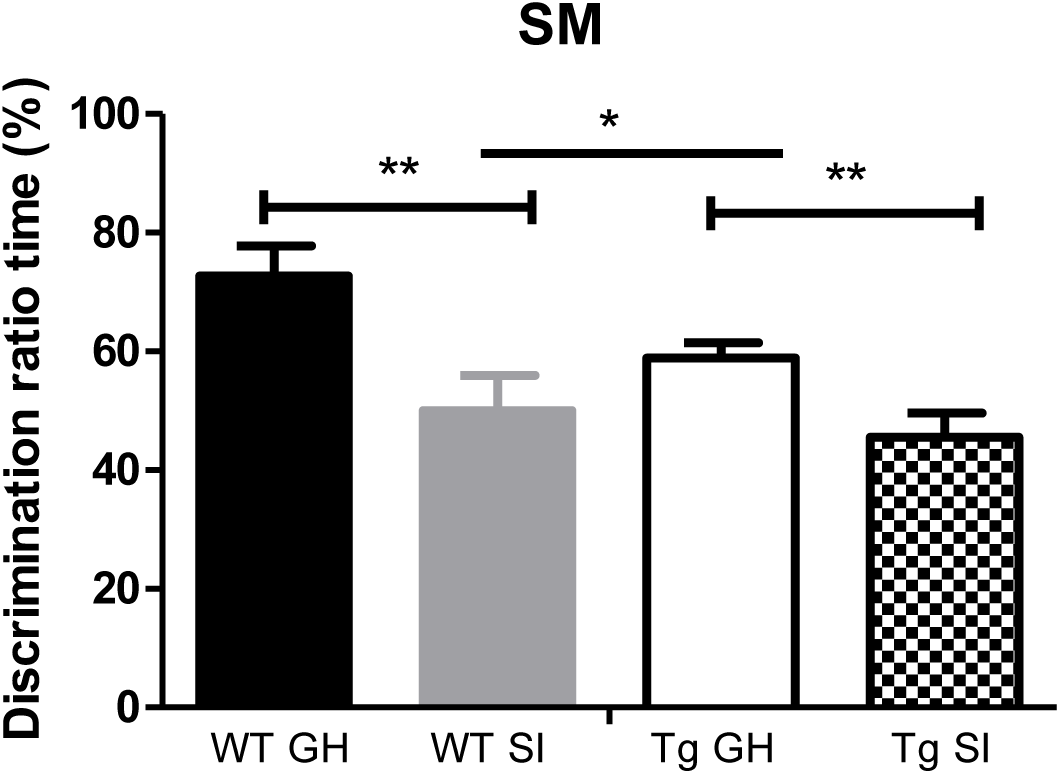
Discrimination rate (DR) of time spent exploring the novel mouse in the social recognition test revealed a difference in social retention memory. Time spent near the stranger mouse in the social memory test revealed two main effects. Firstly, a main effect of genotype was found: AD mice explored less the novel stranger mouse compared to WT mice (* *p* < .05). Interestingly, an additional strong effect of social isolation in the time spent exploring the stranger mouse was also found: isolated mice spent less time with the novel stranger compared to group-housed mice (** *p* < .05).

### SI improved fear memory in APP/PS1 mice, but decreased it in WT mice

When looking at the latency to enter the dark chamber on the test trial, a significant main effect of genotype, *F*(1,56) = 2.284, *p* = .015, and housing condition, *F*(1,56) = 4.870, *p* = .035 was found. Also a Genotype x Housing Condition interaction effect was present: *F*(1,56) = 3.675, *p* = .024. No effect for gender, *F*(1,56) = 2.507, *p*=.119 was found. A closer look at the interaction effect revealed that APP/PS1 mice remembered the shock they received the day before better when isolated, for the WT it was an opposite effect: they performed better when group-housed (Figure 8).

**Figure 8.**
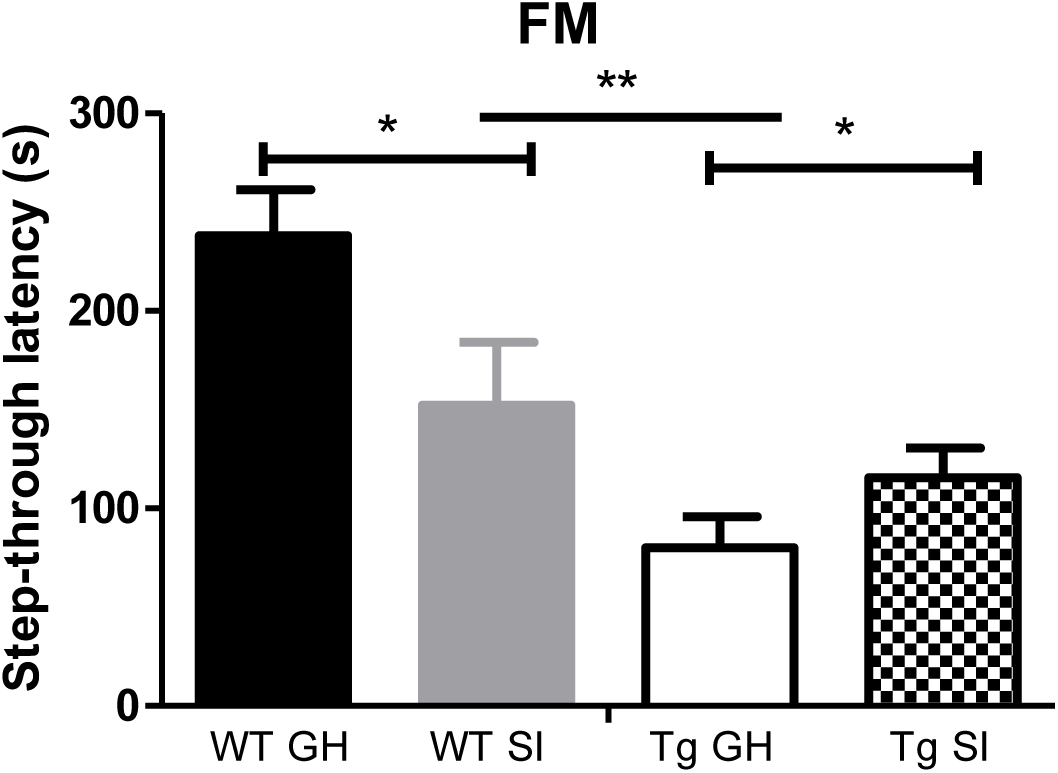
Step-through latencies revealed interesting differences in fear retention memory related to interactions between genotype with housing condition. WT mice were found to perform better when group-housed instead of isolated, however, for APP/PS1 mice this was the opposite: they remembered the shock more when isolation-housed instead of group-housed (** *p* < 0.01; \**p* < 0.05).

## Discussion

Given the ageing population it is of high urgency that the risk factors of AD are known. Both perceived and objective social isolation have long been known as factors that impact overall health, and now it is important to take a closer look at the relationship between social isolation and AD. The aim of this study was to investigate the effects of SI on the development of AD symptomatology by testing both APP/PS1 and healthy WT mice. Part of the animals was socially isolated by housing them individually, while others were group housed. The animals were about 14 months of age when the testing took place and were thus middle-aged. A specific behavioural test battery was used to examine the effects of SI on cognitive functioning, emotionality and sociability.

Three tests were used as a measure of exploration, anxiety and depression-like behaviours: the EPM, the OF and the FST. In both the EPM and OF, APP/PS1 animals were found to exhibit significantly less exploration than their WT counterparts, as measured with proportion of visits to the open arms and time spent in the centre respectively. Therefore we can infer that Tg mice were more anxious and avoidant. No main or interaction effects were found of housing conditions, so GH and SI mice were equally explorative. The FST, which assesses depressive symptoms and despair rather than exploration and anxiety, also only revealed an effect of genotype. APP/PS1 mice were found to be more prone to a depressive-like state than WT mice. Isolation did not increase this in Tgs or install a difference in WTs.

Sociability or social exploration was assessed using the SE test. APP/PS1 mice were found to spend significantly less time with the stranger congener by remaining in the periphery and significantly less time exploring the stranger congener than WT mice. This is in line with previous reports of deficits in sociability in APP/PS1 mice (Filali et al., 2011). Translating animal findings to complex human social behaviours such as empathy should be done with caution. The findings are more easily translatable to social withdrawal and communicative deficits. For example, it has been found that subjects spend increasingly less time interacting with others as dementia progresses (Van der Jeugd et al., 2016). Surprisingly, no significant effects of housing conditions were found, possibly meaning that social exploration has an essential drive or motivation in social species, whether housed separate or not.

The Y-maze was used to assess working memory deficits. The Y-maze revealed no evidence for a difference between groups, however, a trend towards lower memory performance in APP/PS1 mice was present. This was not influenced by housing condition. While the Y-maze is a test of short-term memory, the NOR tap long-term memory processes. When looking at the DR of visits to the novel object, that is, by taking into account the total number of visits to both the novel and the familiar object, WT mice were found to discriminate the novel object better from the familiar one compared to APP/PS1 mice. Housing did not additionally affect this genotype difference. These findings are surprising given that Huang et al. (2011; 2015) found that SI exacerbates both short- and long-term spatial memory deficits in APP/PS1 mice. However, the mice used in their study were old rather than middle aged and spatial memory was assessed using the Y-maze and the Morris water maze.

However, when looking at another form of memory, social recognition memory, we found that APP/PS1 mice had poorer social retention memory compared to WT mice. Moreover, social isolation on itself impacted social retention memory, both WT and APP/PS1 mice that were isolated performed worse compared to their group-housed conspecifics. This points towards the detrimental effects of adult social isolation, in both health and disease.

This is further supported by the fear memory test results, where SI reduces associative memory processing in healthy WT mice. The main psychological symptom observed in AD is indeed memory impairment (like in object and social memory tests), but now we speculate that recall may be influenced when an emotional component is associated with a (negative) event. In AD subjects this can even be enhanced through social isolation, as our (single) foot-shock experiment show. According to our findings, isolated housing is a risk factor for memory dysfunctions. It can be concluded that results concerning the effects of SI on memory are not equivocal. While working memory and long-term object memory is unaffected by SI, there seems to be an impact on short-term social memory and long-term fear memory. This needs further research into the mechanisms of memory formation in healthy and disease, and in loneliness as well as ‘normal’ social settings.

We can conclude, that if social isolation is a risk factor for AD, it therefore could be a modifiable one. Targeting isolated individuals of all ages may therefore have the potential to decrease AD in the population, which is currently one of the ultimate priorities of our global public health. Also the possibility that re-socialization after a period SI is able to undo any negative effects demands further investigation. If SI has a deleterious impact on cognition, re-socialization, physical or cognitive stimulation could have the potential of functioning as a protective treatment of AD.

